# Mass spectrometry imaging of phosphatidylcholine metabolism in lungs administered with therapeutic surfactants and isotopic tracers

**DOI:** 10.1101/2020.10.07.330530

**Authors:** Shane R. Ellis, Emily Hall, Madhuriben Panchal, Bryn Flinders, Jens Madsen, Grielof Koster, Ron. M. A. Heeren, Howard W. Clark, Anthony D. Postle

**Author notes:** Hair Diagnostix, Dutch Screening Group, Gaetano Martinolaan 63A, 6229 GS Maastricht, The Netherlands.

## Abstract

Mass spectrometry imaging (MSI) visualises molecular distributions throughout tissues but is blind to dynamic metabolic processes. Here, MSI with high mass resolution together with multiple stable isotope labelling provided spatial analyses of phosphatidylcholine (PC) metabolism in mouse lungs. Dysregulated surfactant metabolism is central to many respiratory diseases. Metabolism and turnover of therapeutic pulmonary surfactants were imaged from distributions of intact and metabolic products of an added tracer, universally ^13^C-labelled dipalmitoyl PC (U[^13^C]DPPC). The parenchymal distributions of newly synthesised PC species were also imaged from incorporations of *methyl*-D_9_-choline. This dual labelling strategy demonstrated both lack of inhibition of endogenous PC synthesis by exogenous surfactant and location of acyl chain remodelling processes acting on the U[^13^C]DPPC-labelled surfactant, leading to formation of polyunsaturated PC lipids. This ability to visualise discrete metabolic events will greatly enhance our understanding of lipid metabolism in diverse tissues, and has potential application to both clinical and experimental studies.

## Introduction

Phospholipids represent the major surface-active component of lung surfactant. The phospholipid content of surfactant contains high levels of the disaturated species dipalmitoylphosphatidylcholine (DPPC; PC16:0/16:0) that is maintained by a complex intra-alveolar metabolism. Unsaturated PC lipids synthesised on the endoplasmic reticulum are subject to acyl remodelling catalysed by sequential phospholipase A_2_ and lysoPC acyltransferase (LPCAT) activities^1, 2^, intracellular trafficking into lamellar body storage vesicles and secretion into the alveolus to form the mature DPPC-enriched surfactant. Surfactant enriched in DPPC is then secreted into the alveolar lining fluid by exocytosis of lamellar bodies, followed by rapid adsorption to the air:liquid interface. Surfactant is subsequently catabolised by ATII cell endocytosis^3^, metabolism by alveolar macrophages^4^ and loss up the bronchial tree^5^. A proportion of the surfactant taken up by ATII cells is subsequently recycled into lamellar bodies for re-secretion into the alveolus^6, 7^.

Prior work by some of us used the incorporation of deuterated *methyl*-D_9_-choline into PC molecular species to probe surfactant metabolism in greater detail from biological extracts^8, 9^. This work characterised the mechanisms of acyl remodelling in surfactant PC synthesis by animal models^9^, adult volunteers^10^ and acute respiratory disease syndrome (ARDS) patients^11^. Recently an alternative acyl remodelling mechanism involved in surfactant metabolism was demonstrated, whereby a proportion of surfactant DPPC is selectively remodelled into PC species containing the polyunsaturated fatty acids arachidonate (20:4) and docosahexaenoate (22:6)^12^. This study administered exogenous therapeutic surfactants to adult mice, with tracer amounts of the universally ^13^C-labelled (U^13^C) isotopomer of DPPC (U^13^C-DPPC) added to the surfactant. In addition to quantifying overall surfactant catabolism, this experimental approach enabled characterisation of recycling mechanisms of surfactant DPPC by following the incorporations into intact PC species of U^13^C-labelled fragments derived from U^13^C-DPPC hydrolysis.

Lung surfactant components such as DPPC can also be analysed by mass spectrometry imaging (MSI), which offers a powerful approach to study the distributions of lipids throughout biological tissues^13, 14^. Matrix-assisted laser desorption/ionisation (MALDI)-MSI has demonstrated enriched location of polyunsaturated PC species within the bronchial tree^15^, a distribution potentially linked to the involvement of oxidation products of arachidonate in the pathology of inflammatory lung diseases^16, 17^. However, the location of synthesis within lung tissue of PUFA-containing PC species, both directly by the CDP:choline pathway and by acyl remodelling, has not been established. This information is lacking because conventional MSI is only capable of providing static lipid compositions within a given tissue region at a fixed moment in time. This static limitation can be overcome by coupling MSI with stable isotope labelling: isotope uptake into lipid metabolic processes can be detected in a time-resolved manner, reflected by measurements of mass-shifted lipid signals containing the isotopic tracer or the products of its metabolic conversion. Such approaches have been more widely applied using secondary ion mass spectrometry (SIMS), but due to the extensive fragmentation observed by SIMS whereby often only elemental or diatomic fragments are observed (e.g., Nano SIMS), it is difficult to study the metabolism of individual lipid species^18, 19^. Recently several groups have coupled isotope labelling with MALDI^20^ or with desorption electrospray ionisation (DESI)^21^, both of which enable detection of intact lipid species. These studies utilised heavy water labelling of mice to study region-specific lipid synthesis in breast cancer and mouse brain tissue, respectively. The widespread and non-specific nature of heavy water labelling were significant limitations of these studies. These limitations, together with the modest mass resolving power of the MSI instrumentation used, meant many lipid signals, representing combinations of different labelled and unlabelled species, and different metabolic processes, remained unresolved.

Here we used MALDI-MSI at ultra-high mass resolving power coupled a dual labelling approach utilising both *methyl*-D_9_ choline chloride and U^13^C-DPPC administered to mice to visualise lung and surfactant lipid metabolism. We report the simultaneous characterisation and visualisation of surfactant PC synthesis and catabolism throughout the lung tissue.

## Results

### MSI of isotopically labelled lipids

We aimed to analyse lipid metabolism in mouse lung tissue after dosing intraperitoneally with D_9_-choline chloride and intranasally with an exogenous therapeutic surfactant containing U-^13^C-DPPC. Both the complex mass spectra generated by MALDI analysis of biological tissue and the expected low abundance of labelled lipid species (~1–2% relative to the corresponding unlabelled lipid species) make the unambiguous detection of both D_9_-choline–labelled PC and U-^13^C-DPPC directly from lung tissue a challenge. Here we have utilised the high resolving power of an Orbitrap mass spectrometer to enable unambiguous detection of isotopically labelled species in the presence of isobaric interferences arising from endogenous lipids. The average mass spectrum of a mouse lung tissue showed that protonated, sodiated and potassiated ions of PC lipids dominate the spectrum with PC32:0 (predominately PC16:0/16:0), the major lipid component of lung surfactant, providing the highest signal intensities with the base peak at *m/z* 756.55134 assigned as the [PC32:0+Na]^+^ ion (Fig. 1a). Importantly, we detected D_9_-labeled PC32:0, corresponding to PC32:0 synthesised over the 12 h between label injection and sacrifice, with the [D_9_-PC32:0+Na]^+^ ion detected at *m/z* 765.60785 with an intensity of ~1% compared to the unlabelled variant (Fig. 1b). The corresponding protonated and potassiated ions were also observed (Supplementary Fig. 1). We detected [U^13^C-PC32:0+Na]^+^ at *m/z* 796.68538 that arises exclusively from the therapeutic surfactant (Fig. 1c), as well as its corresponding protonated and potassiated ions (Supplementary Fig. 2). Identification of both labelled PC species ions was further corroborated by tandem mass spectrometry (MS/MS) with specific fragments consistent with the expected deuterium ([D_9_-PC32:0+Na]^+^, Fig. 1d) and ^13^C ([U^13^C-PC32:0+Na]^+^, Fig. 1e) labelling patterns detected with high mass accuracy (< 1 ppm mass error). These results confirm the ability to detect both newly synthesised PC lipids via the Kennedy pathway and D_9_-choline incorporation as well as exogenous surfactant using U^13^C-PC32:0 as a specific marker ion directly from lung tissue sections using MALDI-MSI.

**Figure 1.**
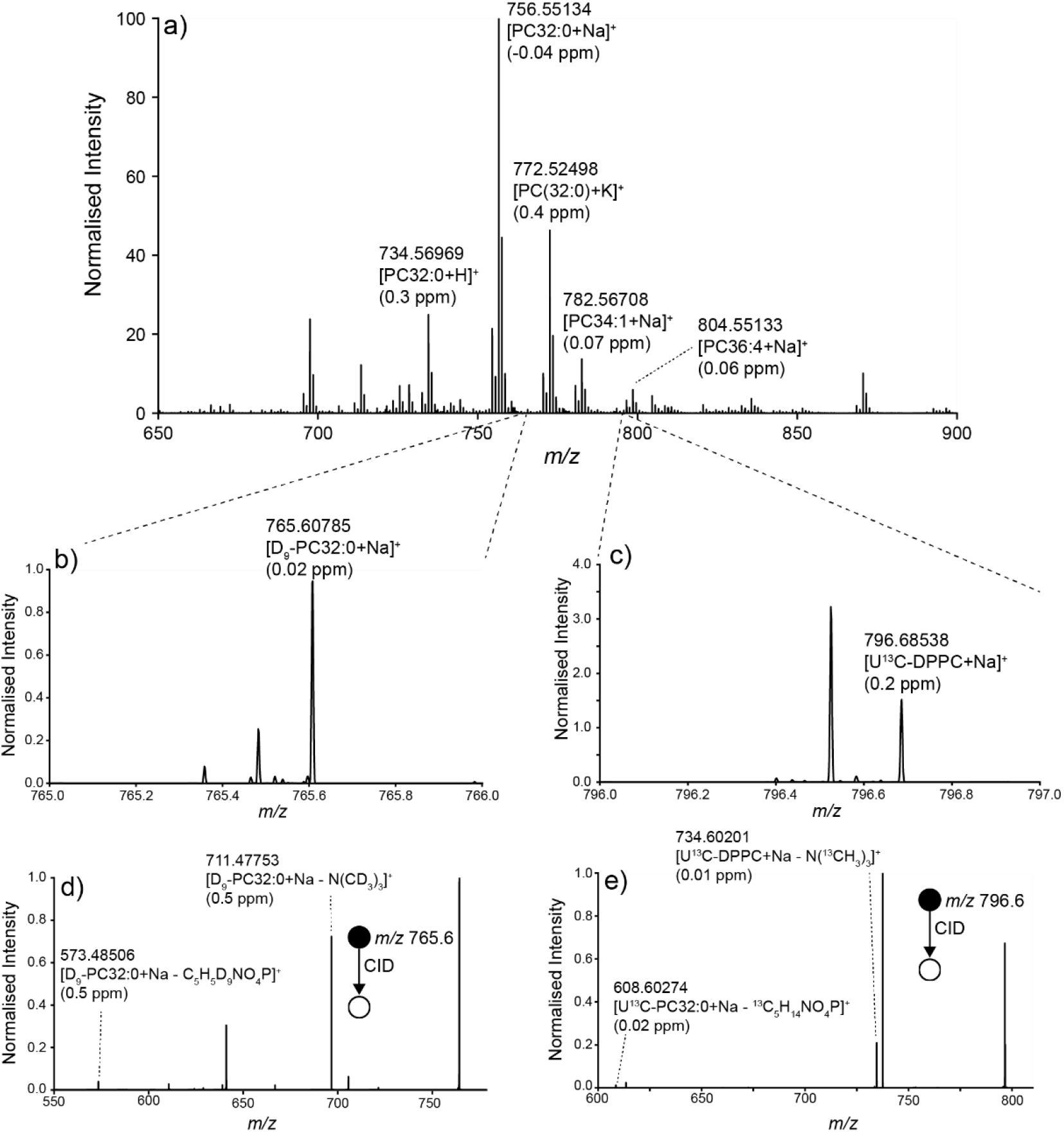
(a) Averaged positive-ion MALDI spectrum of mouse lung tissue administered with D_9_-choline and U^13^C-DPPC-containing CHF5633 surfactant 12 h prior to sacrifice. Lipid identifications and parts-per-million (ppm) mass errors for several abundant lipid species are indicated. (b, c) Zoom-in of the spectrum in (a) at the *m/z* 765–766 range (b) demonstrating the detection of [D_9_-PC32:0+Na]^+^ and at the *m/z* 796–797 range (c) demonstrating the detection of [U^13^C-DPPC+Na]^+^. (d, e) MS/MS spectra for precursor ions at *m/z* 765.5 ± 0.5 (d) and *m/z* 796.6 ± 0.5 (e) corresponding to [D_9_-PC32:0+Na]^+^ and [U^13^C-DPPC+Na]^+^, respectively, with supporting fragments annotated. Additional peaks in the MS/MS spectra arise from the co-isolation and fragmentation of isobaric ions. DPPC = PC16:0/16:0.

### Imaging of surfactant phosphatidylcholine metabolism in mouse lungs

Next, we sought to study the distribution of both D_9_-choline–labelled PC32:0 and intranasally dosed exogenous surfactant within mice lung tissue. We administered two different surfactants - synthetic CHF5633 and porcine lung-derived Poractant alfa. These preparations have different phospholipids, with CHF5633 being composed of a 50/50 mixture of PC16:0/16:0 and PG16:0/18:1 while Poractant alfa contains the full complement of surfactant lipids. U^13^C-PC16:0/16:0 was added to both surfactants to monitor exogenous surfactant metabolism before intranasal administration to mice that were also injected with *methyl*-D_9_-choline chloride to assess endogenous lung PC synthesis occurring via the Kennedy pathway^22^.

### CHF5633 synthetic surfactant

The distribution of several sodiated PC lipid ions throughout the mouse lung 12 h after administration of CHF5633 are shown in Figure 2. The optical image of the haematoxylin and eosin (H&E) stained of the same tissue after MSI is shown in Figure 2a. The U^13^C-DPPC ion at *m/z* [M+Na]^+^ 796.6855 was located in one parenchymal area but not within bronchioles, demonstrating one extreme example of the regional deposition (Fig. 2b). As expected, the distribution of unlabelled DPPC at *m/z* [M+Na]^+^ 756.5514 (Fig. 2c) partially mirrored that of U^13^C-DPPC (Fig. 2a) due to the excess amount of unlabelled DPPC in the CHF6533 surfactant. By contrast, the distribution of ion signals corresponding to the incorporation *methyl*-D_9_-choline into PC32:0 via the Kennedy pathway (Fig. 2d; [M+Na]^+^, *m/z* 765.6079) exhibited a reduced intensity in regions where U^13^C-DPPC was highest, demonstrated by the ratio of D_9_-PC32:0+Na^+^/PC32:0+Na^+^ (Fig. 2e); this observation could suggest inhibition of endogenous PC synthesis by the exogenous surfactant. However, both unlabelled PC32:1 (Fig. 2f, [M+Na]^+^, *m/z* 754.5359), which is an integral component of mouse lung surfactant but absent from CHF5633, and D_9_-PC32:1 (Fig. 2g, [M+Na]^+^, *m/z* 763.5923) were similarly present at lower intensity in regions enriched in CHF5633. Consequently, the ratio of [D_9_-PC32:1+Na]^+^/[PC32:1+Na]^+^ was more uniform across the lung parenchyma (Fig. 2h) with no decrease in areas of high U^13^C-DPPC signal. Similar distributions were apparent for other endogenous PC species with a parenchymal distribution and were consistent for all sections analysed (data not shown). These results suggest the decreased D_9_-[PC32:0+Na]^+^/[PC32:0+Na]^+^ ratios are a consequence of the regional, non-specific accumulation of the DPPC-rich CHF6533 surfactant, rather than inhibition of PC synthesis. This is supported by extracted region-specific spectra that show regions of high U^13^C-DPPC to contain 2–3 fold higher signal for unlabelled DPPC than surrounding regions (Supplementary Fig. 3). Although some lipid signal was observed on the glass slide adjacent to the tissue sections, which may arise due to smearing artefacts during tissue mounting, this did not alter the on-tissue distributions.

**Figure 2.**
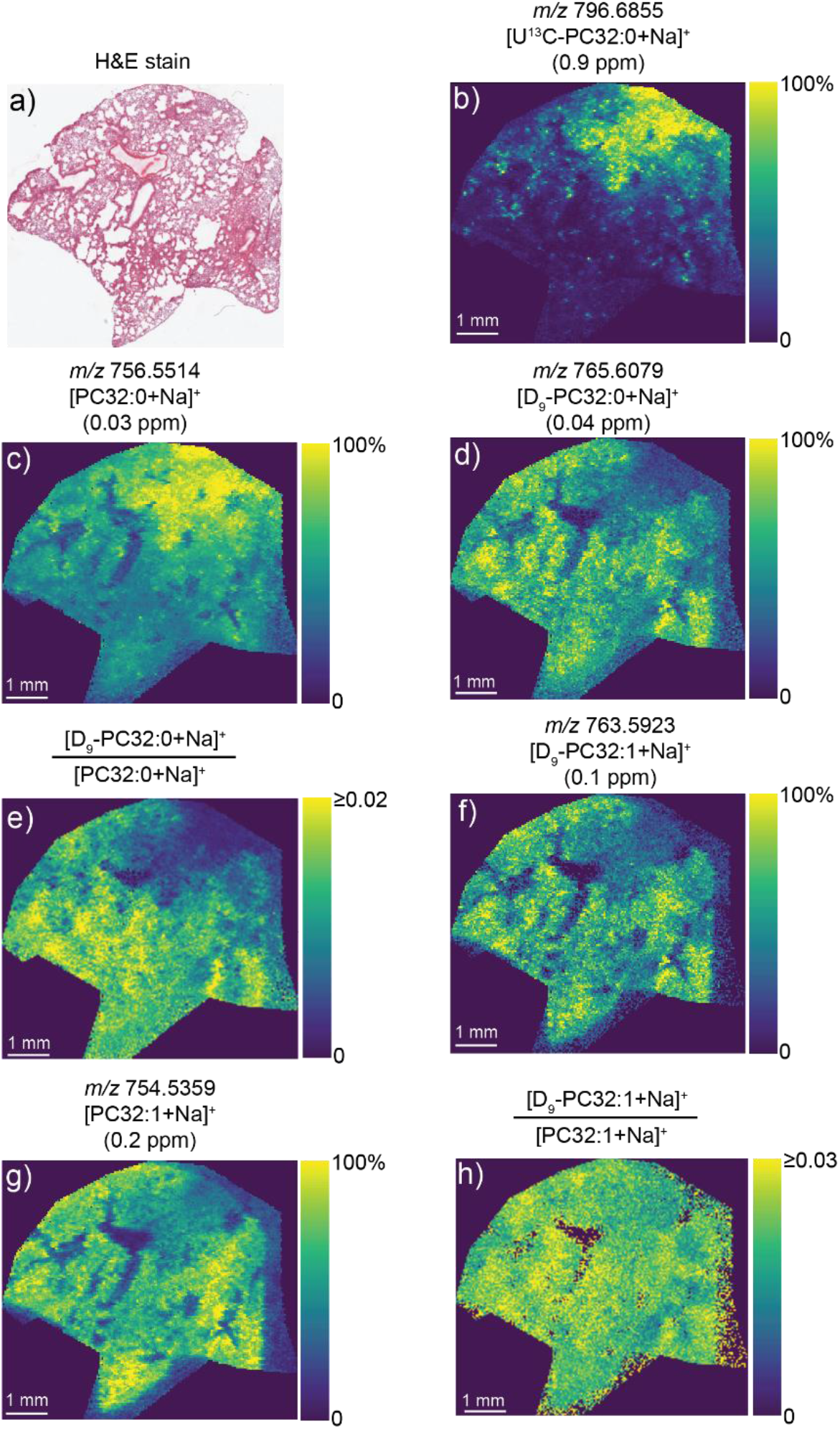
MALDI-MSI data from the same mouse lung tissue analysed in Figure 1. (a) Optical image of the post-MSI, H&E-stained tissue section. (b–d, f–g) Ion images of (b) *m/z* 796.6855 ([U^13^C-DPPC+Na]^+^), (c) *m/z* 756.5514 ([PC32:0+Na]^+^, (d) *m/z* 765.6079 ([D_9_-PC32:0+Na]^+^, (f) *m/z* 763.5923 ([D_9_-PC32:1+Na]^+^) and (g) *m/z* 754.5359 ([PC32:1+Na]^+^). (e, h) Ratio images of (e) [D_9_-PC32:0+Na]^+^:[PC32:0+Na]^+^ and (h) [D_9_-PC32:1+Na]^+^:[PC32:1+Na]^+^. Part-per-million (ppm) mass errors are indicated in parentheses. All images were visualised using total-ion-current normalisation and using hotspot removal (high quantile = 99%). DPPC = PC16:0/16:0.

In agreement with previous reports^15^, MSI demonstrated spatial segregation of unlabelled PC species, with the polyunsaturated PC species PC36:4 (Fig. 3a, [M+Na]^+^, *m/z* 804.5514) and PC38:6 (Fig. 3b, [M+Na]^+^, *m/z* 828.5514) restricted to bronchiolar regions (Figure 3c–d). Similar results were obtained obtained from another animal sacrificed 6 h after label administration (Supplementary Fig. 4). Little to no signal was observed for exogenous CHF5633-specific U^13^C-DPPC in the bronchiolar regions. Unfortunately, due to the low abundance of these polyunsaturated species compared to PC32:0 and PC32:1 that form major lipid components of surfactant, combined with the presence of an unresolved isotopologue from another lipid-related ion, the corresponding D_9_-containing PC36:4 and PC38:6 ions could not be unambiguously detected.

**Figure 3.**
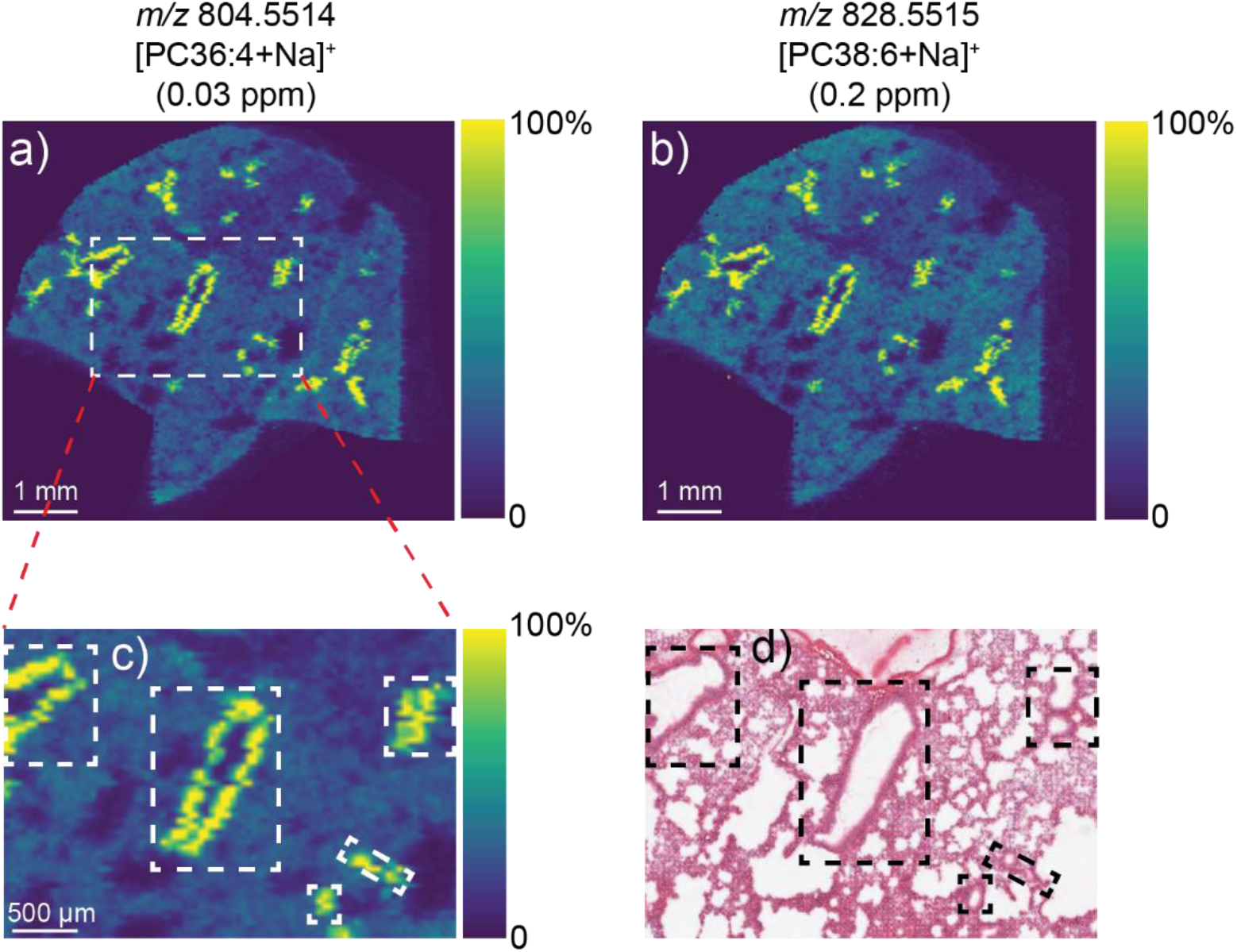
MALDI-MSI data from the same mouse lung tissue analysed in Figure 1. Ion images of (a) *m/z* 804.5514 ([PC36:4+Na]^+^) and (b) *m/z* 828.5515 ([PC38:6+Na]^+^). (c) Enlarged image of the box in (a) showing these polyunsaturated lipids localised to the bronchiolar regions, as identified by the post-MSI H&E-stained section (d). All images were visualised using total-ion-current normalisation and using hotspot removal (high quantile = 99%).

Incorporation of *methyl*-D_9_-choline also facilitated analysis of the distribution patterns of other newly synthesised PC species. Other PC species characteristic of lung tissue rather than surfactant, such as PC34:1 and PC34:2^23, 24^ were distributed more equally between bronchiolar and parenchymal regions both for unlabelled and D_9_-choline-labelled species (Supplementary Fig. 5). Consistent distributions as those described above were observed from lung tissue obtained from other dosed animals (Supplementary Fig. S6). For all labelled ions discussed above, no signal was observed in a control lung tissue that was not administered with either D_9_-choline or U^13^C-DPPC (Supplementary Fig. 7).

### Poractant alfa porcine surfactant

Nasal administration of the porcine-derived Poractant alfa surfactant typically achieved a more widespread distribution in the lungs compared with CHF5633, illustrated by the distribution of the [U^13^C-DPPC+Na]^+^ ion at 12 h post-dosing (Fig. 4a). Analysis of another mouse 18 h after administration showed comparable results (Supplementary Fig. 8). We observed a more uniform distribution of unlabelled [PC32:0+Na]^+^ (Fig. 4b) and [D_9_-PC32:0+Na]^+^ (Fig. 4c and 4d), although the normalised signal intensity for [D_9_-PC32:0+Na]^+^ was still slightly more intense in lung regions with a relatively low U^13^C-DPPC signal due to the presence of unlabelled PC32:0 in the Poractant alfa surfactant.

**Figure 4.**
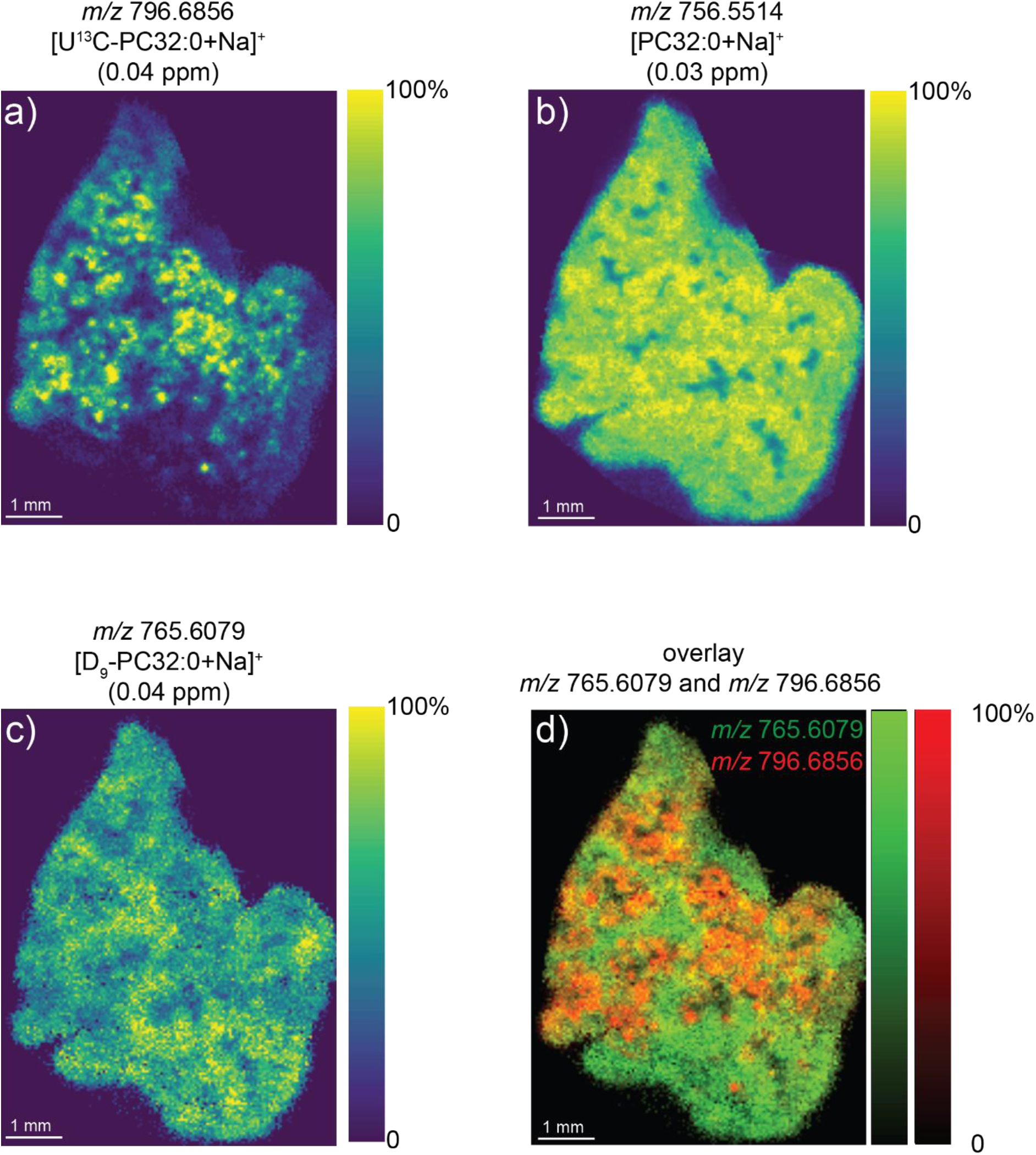
MALDI-MSI data of mouse lung tissue after administration with D_9_-choline and U^13^C-DPPC–containing Poractant alfa surfactant (labels administered 12 h prior to sacrifice). Ion images of (a) *m/z* 796.6856 ([U-^13^C-DPPC+Na]^+^), (b) *m/z* 756.5154 [PC32:0+Na]^+^ and (c) *m/z* 765.6079 ([D_9_-PC32:0+Na]^+^). (d) Overlay image of D_9_-PC32:0+Na]^+^ (red) and [U^13^C-PC32:0+Na]^+^ (green). Part-per-million (ppm) mass errors are indicated in parentheses. All images were visualised using total-ion-current normalisation and using hotspot removal (high quantile = 99%). DPPC = PC16:0/16:0.

### Alveolar metabolism of U^13^C-DPPC–labelled surfactant *in vivo*

Previous analysis of the *in vivo* metabolism of U^13^C-DPPC–labelled surfactants in mice by shotgun lipidomics analysis of extracts identified the labelled ^13^C_5_-choline and ^13^C_24_-lysoPC16:0 moieties, generated by the respective phospholipase D and phospholipase A_1_/A_2_ enzymatic hydrolysis of U^13^C-DPPC, and their incorporation into PC molecular species^12^. The molecular specificity for ^13^C_5_-PC lipid synthesis closely mirrored that observed for D_9_-choline labelling. This indicates that ^13^C_5_-choline generated by U^13^C-DPPC hydrolysis equilibrated with endogenous unlabelled choline and was then incorporated into PC by the CDP:choline (Kennedy) pathway^12^. By contrast, re-acylation of ^13^C_24_-LPC16:0 in both lung tissue and bronchoalveolar lavage was much more restricted, generating primarily only two polyunsaturated PC species, ^13^C_24_-PC16:0_20:4 and ^13^C_24_-PC16:0_22:6^12^. Our analysis of the lavage cell pellet from this study^12^ showed a higher enrichment of both these species than in either lung tissue or bronchoalveolar lavage (Supplementary Fig. 9). As this cell pellet contained both alveolar macrophages and the more easily precipitated lung surfactant, we hypothesised that re-acylation of ^13^C_24_-LPC16:0 takes place in alveolar macrophages. To test this, mice received nasal administration of U^13^C-DPPC–labelled CHF5633, an intraperitoneal injection of *methyl*-D_9_-choline chloride (1 mg) 18 hours later, were euthanized at 21 hours after the first administration and lung tissue taken for analysis. Alveolar macrophages were purified by centrifugation at 400 × *g* for 10 min, followed by differential adherence and washing to remove surfactant. There was negligible incorporation of the D_9_-choline label into alveolar macrophage PC, indicating a very low rate of PC synthesis *de novo* (data not shown). This contrasts strikingly with the high rate of surfactant PC synthesis by alveolar type II epithelial cells, which was greatest at 3 h^12^. In comparison, active re-acylation of ^13^C_24_-LPC16:0 via the Lands cycle^23, 24^ by alveolar macrophages *in vivo* could be readily demonstrated 21 hours after CHF5633 administration^12^. Increased unlabelled PC32:0 in alveolar macrophages showed the uptake of exogenous surfactant by these cells (Fig. 5a). However, analysis of ^13^C_24_-PC also showed considerable metabolism of this intracellular exogenous surfactant over the same time period. The relative abundance of U^13^C-DPPC decreased from the initial administered value of 100% at 0 hours, while those of ^13^C_24_-PC16:0_20:4 and ^13^C_24_-PC16:0_22:6 re-acylation products increased (Fig. 5b). This finding was observed for all 8 treated mice, with the mean relative abundances of U^13^C-DPPC, ^13^C_24_-PC16:0_20:4 and ^13^C_24_-PC16:0_22:6 being 38.7 ± 8.9%, 36.8 ± 9.7% and 24.5 ± 6.4% (mean ± s.d.) relative to U^13^C-DPPC at t=0 (data not shown). There was no detectable synthesis of ^13^C_24_-DPPC by alveolar macrophages *in vivo*, which is consistent with the absence of DPPC synthesis by acyl remodelling in this cell type^25^.

**Figure 5.**
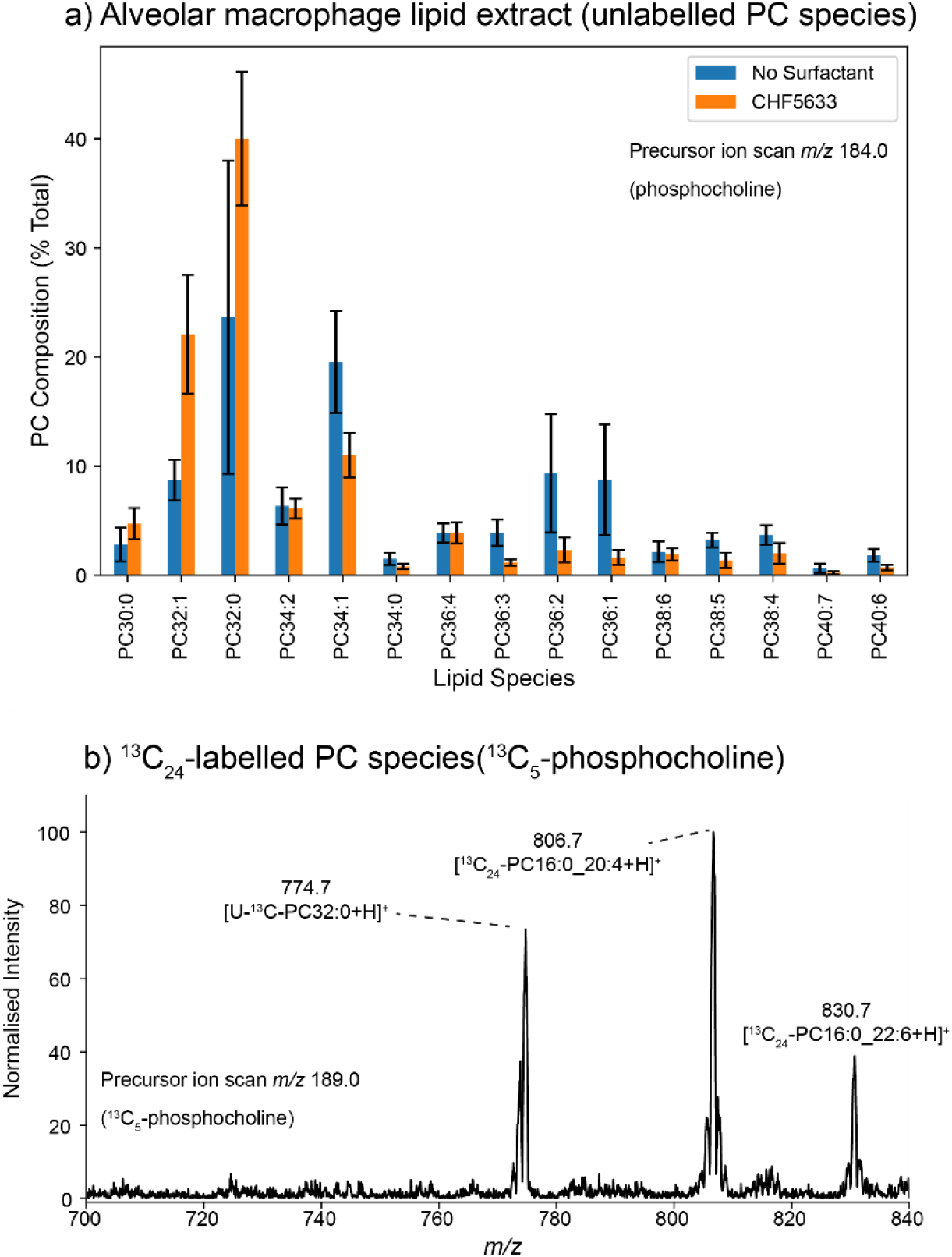
Alveolar macrophage metabolism of labelled exogenous surfactant. U^13^C-DPPC– labelled CHF5633 surfactant (4 mg in 50 μl) was administered intranasally to mice, followed 18 h later by an intraperitoneal injection with *methyl*-D_9_-choline chloride (1 mg). Mice were sacrificed 3 h later, their lungs were lavaged and alveolar macrophages isolated by differential adherence. After washing with saline, macrophage lipids were extracted and analysed by ESI-MS/MS. (a) Unlabelled PC lipid composition of isolated alveolar macrophages using a precursor ion scan of *m/*z 184.0. Orange bars correspond to mice administered with CHF5633 surfactant and blue bars mice not administered with any exogenous surfactant. (b) Precursor ions scan for *m/*z 189.0 for detection of PC species containing the 5^13^C phosphocholine headgroup derived from U^13^C-DPPC. Error bars correspond to ± 1 standard deviations (*n* = 8 per group). DPPC = PC16:0/16:0.

### Imaging surfactant acyl remodelling *in vivo*

If re-acylation of ^13^C_24_-LPC16:0 is restricted to alveolar macrophages *in vivo*, then ^13^C_24_-PC16:0_20:4 should not co-localise with unlabelled PC36:4 in the bronchioles. Consequently, we monitored the distribution of ^13^C_24_ PC species in mouse lungs 12 and 18 h after administration of U^13^C-DPPC-labelled Poractant alfa, specifically chosen for this analysis rather than CHF5633 for two reasons. Firstly, the more uniform distribution of the porcine surfactant (Fig. 4) facilitated a clearer analysis of the distribution of surfactant metabolites. Secondly, enrichment of U^13^C-DPPC was higher in administered Poractant alfa than in CHF5633^12^, thereby providing sufficient signal intensity for imaging of the low abundance ^13^C_24_-labelled acyl chain remodelling products.

In our MALDI-MSI analysis we detected the [^13^C_24_-PC16:0_20:4+Na]^+^ ion at *m/z* 828.6320 (Fig. 6a) with a relative intensity of ~0.1% compared to the [DPPC+Na]^+^ ion in the mean spectrum. This compound could be assigned with high confidence due to the low mass error of 0.2 ppm and absence of this signal in an undosed mouse lung, in spite of its low abundance (Supplementary Fig. 7). Further confirmation for the assignment of this ion as [^13^C_24_-PC16:0_20:4+Na]^+^ was provided by the MS/MS spectrum (Fig. 6b) that revealed fragment ions at *m/z* 766.54786 and 640.54869, corresponding to the neutral losses of N(^13^CH_3_)_3_ (−0.6 ppm mass error) and ^13^C_5_H_15_NO_4_P (−0.6 ppm mass error). These fragment ions are unambiguously attributable to a ^13^C-labelled phosphocholine headgroup. Unlabelled [PC36:4+Na]^+^ (green) was observed primarily in the bronchioles of mouse lungs at 12 h (Fig. 6c) and 18 h (Fig. 6d), along with a corresponding distribution of [U-^13^C-DPPC Na]^+^ (blue) and [^13^C_24_-PC16:0_20:4+Na]^+^ (red) that is synthesised by successive de-acylation/re-acylation processes of U^13^C-DPPC through the Lands cycle. These processes led to replacement of a U^13^C-16:0 fatty acyl with an unlabelled 20:4 acyl chain, leading to the formation of ^13^C_24_-PC16:0_20:4. Thus, the detection of both the lipid substrate and end products of this enzymatic-driven conversion provides the first direct observation of acyl remodelling events in tissue using MSI. Signal for [U^13^C-DPPC+Na]^+^ was absent from the bronchioles and distributed heterogeneously throughout the parenchymal area. While this does not allow us to exclude the possibility of PC16:0/16:0 → PC16:0_20:4 metabolism within the bronchioles, the observation of [^13^C_24_-PC16:0_20:4+Na]^+^ provides direct evidence for PC16:0/16:0 → PC16:0_20:4 enzymatic synthesis within the lung parenchyma. The absence of ^13^C_24_-PC16:0_20:4 and U^13^C-DPPC in the bronchial regions demonstrated the lack of surfactant uptake and PC metabolism by the bronchial epithelium. We found reasonable but not complete co-localisation between ^13^C_24_-PC16:0_20:4 and U^13^C-DPPC, a perhaps not surprising observation as alveolar surfactant and macrophages had been removed by lavage before the lungs were frozen. These distributions then presumably represent the patterns of cellular uptake of surfactant by ATII cells and macrophages as well as that of surfactant acyl remodelling in lung tissue-resident alveolar macrophages. Evidence for the 22:6 acyl chain substitution on U^13^C-DPPC was observed but with insufficient signal for imaging. While the [^13^C_24_-PC16:0/22:6+Na]^+^ ion was absent in the full-scan MSI data, MS/MS of expected precursor ion revealed a characteristic neutral loss of N(^13^CH_3_)_3_ thereby providing direct evidence for a [24^13^C-PC16:0/22:6+Na] ion (Supplementary Fig. 10). For enhanced sensitivity, this MS/MS data were obtained following accumulation of precursor ions over a larger sample area before fragmentation and detection. The results thereby present evidence for the specific acyl remodelling of surfactant DPPC into polyunsaturated PC species by alveolar macrophages *in vivo*.

**Figure 6.**
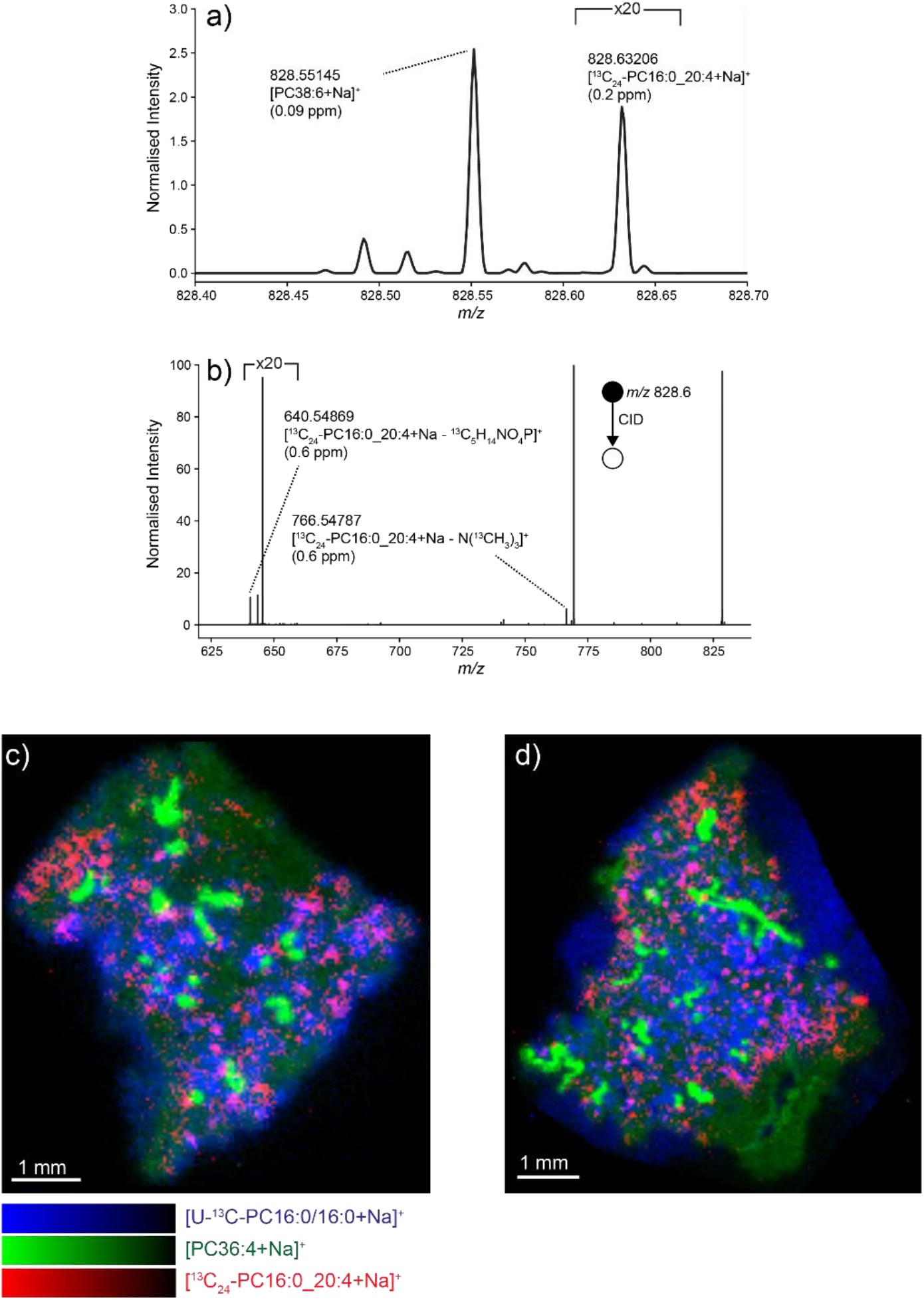
MALDI-MSI of U^13^C-PC16:0/16:0 acyl chain remodelling. (a) Averaged MALDI mass spectrum from lung tissue collected from mice sacrificed 12 h after administration of D_9_-choline and U^13^C-DPPC–containing Poractant alfa surfactant. The ion at *m/z* 828.63206 is assigned as the [M+Na]^+^ ion of ^13^C_24_-PC16:0_20:4 formed by acyl remodelling of U^13^C-PC16:0/16:0 (b) MS/MS spectrum of precursor ions at *m/z* 828.5 ± 0.5 with fragment ions consistent with the presence of [^13^C_24_-PC16:0_20:4+Na]^+^ annotated. Part-per-million (ppm) mass errors are provided in parentheses. (c, d) MALDI-MSI data of [U^13^C-DPPC+Na]^+^ (blue), [PC36:4+Na]^+^ (green) and [^13^C_24_-PC16:0_20:4+Na]^+^ (red) in lung tissue collected from mice 12 h (c) and 18 h (d) after label administration. All images were visualised using total-ion-current normalisation and using hotspot removal (high quantile = 99%).

## Discussion

For the first time we have combined isotopic labelling with MALDI-MSI at high mass resolving power to visualise: (i) uptake and distribution of therapeutic surfactants used to treat a variety of respiratory diseases; and (ii) the location of lipid synthesis and metabolism events throughout biological tissue. Injection of D_9_-choline into the mice enabled the detection and localisation of newly synthesised PC lipids produce by the CDP:choline pathway by virtue of their expected *m/z* shift of 9.0654 (Δ_9D-9H_). These metabolic events were found to be relatively homogenously distributed throughout the mouse lung tissue. The use of a U^13^C-DPPC label not only enabled the visualisation of both CHF5633 and Poractant alfa porcine surfactant uptake independent of endogenous surfactant, but also allowed detection of acyl chain remodelling events within the tissue. In particular, de-acylation/re-acylation of U^13^C-DPPC via the Lands cycle was observed, forming ^13^C_24_-PC16:0_20:4 within the parenchyma. We note that while it is often assumed that this process is initiated by PLA_2_ cleavage of the *sn*-2 fatty acyl, analogous PLA_1_ cleavage^26, 27^, leading to the formation of ^13^C_24_-PC20:4/16:0, is also possible. As a result, our approach not only provides lipid distributions, but the localisation of lipid metabolic events within the tissue.

The metabolism of U^13^C-DPPC by alveolar macrophages *in vivo*, summarised in Figure 5, provides strong circumstantial evidence for a major role for this cell type in the generation of ^13^C_24_-PC16:0_20:4 by de-acylation/re-acylation of DPPC. While the active re-acylation of arachidonate into alveolar macrophage phospholipids has been reported previously^28, 29^, the results presented here are the first demonstration of a similar re-acylation that is specific for docosahexaenoate. If re-acylation of arachidonoate is indeed restricted to alveolar macrophages, this could explain the discrepancy of distribution of ^13^C_24_-PC16:0_20:4 and U^13^C-DPPC shown in Figure 6. This may possibly have been due to migration of alveolar macrophages within the lung parenchyma after initial uptake of the exogenous surfactant.

Surfactant catabolism and turnover have previously been studied both *in vivo*^30^ and in isolated ATII cells^31^ and alveolar macrophages^32^, which are thought to be responsible for the turnover of surfactant phospholipid in healthy lung. However, there has been no demonstration of their relative contributions, as opposed to, for instance secretory phospholipase A_2_^33^, to the decreased bronchoalveolar surfactant concentration in severe respiratory diseases such as acute respiratory distress syndrome. In contrast, the extensive accumulation of surfactant phospholipid and proteins in alveolar proteinosis is clearly due to impairment of GM-CSF– dependent alveolar macrophage metabolism^34^. MSI offers considerable potential to probe these mechanisms in increased detail, particularly as the sensitivity continues to improve, enabling the detection of even lower abundance labelled lipids and their metabolites. This detection sensitivity will be especially important if the relative roles of different individual cell types can be visualised, a possibility that is becoming feasible as current research aims to reduce the spatial resolution of MSI to the single-cell level^35, 36^.

## Materials and Methods

### Preparation of labelled surfactants

U^13^C-DPPC was mixed with the porcine-derived Poractant alfa and the synthetic CHF5633 surfactants as described previously^12^. Isotopic enrichments of U^13^C-DPPC in Poractant alfa and in CHF5633 were, respectively, 7.55% and 2.87% of total DPPC.

### Animal labelling and sample preparation procedure

Mouse lungs used in this study were provided from a previous study of exogenous surfactant turnover in mice^12^. Tissues were stored at −80 °C, shipped on dry ice for MALDI-MSI, then stored at −80 °C until sectioning and analysis. Briefly, animal procedures were approved internally by the University of Southampton Animal Welfare and Ethical Review Body and externally by the Home Office, Animals in Science Regulation Unit. Wild-type (C_5_7BL/6) male mice aged between 8 to 12 weeks were used for this study. Randomisation was not relevant in this study. Similarly, blinding was not practicable as the two surfactant used had different phospholipid compositions. Each mouse was instilled intranasally with 50 μl (4 mg, equivalent to 200 mg/kg body weight) of either Poractant alfa or the synthetic surfactant CHF5633, both containing U^13^C-DPPC. At the same time, each mouse also received a 100 μl intraperitoneal injection of *methyl-*D_9_-choline chloride (10 mg/ml in water). After labelling, the mice were sacrificed by carbon dioxide asphyxia at 6, 12, 18 or 24 hours. Bronchoalveolar lavage was performed in situ with 4 × 0.9 ml PBS, and the recovered bronchoalveolar lavage fluid (BALF) aliquots were combined. BALF was centrifuged at 300 × *g* for 10 min at 4 °C to pellet cells, and the supernatants were then transferred to new vials and stored at −80 °C until extraction. Lung parenchyma was quickly dissected from the main bronchi, placed in cryotubes and snap-frozen in liquid nitrogen. Further details are provided in reference 12.

### Alveolar macrophage labelling *in vivo*

For *in vivo* labelling studies, 8 male wild-type C57BL6 mice were instilled with CHF5633 surfactant containing U^13^C-DPPC, then injected with *methyl*-D_9_-choline chloride 18 hours later. After 3 hours, mice were sacrificed and BALF obtained, from which alveolar macrophages were isolated by centrifugation at 400 × *g* for 10 min BALF Macrophages were purified by adherence to plastic tissue-culture dishes and were subsequently washed three times with 0.9% saline to remove adherent surfactant.

### Macrophage phospholipid analysis

Total lipids were extracted by scraping adherent alveolar macrophages into 800 μl of 0.9% saline after adding dimyristoyl PC (10 nmoles) as internal standard, followed by extraction with dichloromethane (2 ml), methanol (2 ml) and water (1 ml)^37^. After mixing and centrifugation at 1500 × *g*, 20 °C for 10 min, the dichloromethane-rich lower phase was recovered, dried under a stream of nitrogen gas and stored at −20 °C until analysis by mass spectrometry. Mass spectrometry analysis was performed on a Waters XEVO TQ-MS instrument using electrospray ionisation as described previously^9^. Unlabelled PC and newly synthesised PC labelled with D_9_-choline were detected using precursor ion scans of the PC headgroup fragment ions at *m/z* +184 and *m/z* +193, respectively. Precursor ion scans of *m/z* +189 detected the PC species containing five labelled ^13^C atoms in their phosphocholine head group.

### Tissue preparation for MALDI-MSI

Frozen lung tissue was sectioned for MALDI-MSI analysis using a Leica CM 1860 UV cryomicrotome (Leica Microsystems, Wetzlar, Germany) at a temperature of −20 °C to produce 12 μm–thick tissue sections, which were thaw mounted onto clean indium tin oxide (ITO)-coated glass slides (Delta Technologies, Loveland, CO, USA). Tissue sections were stored at −80 °C until matrix application and analysis. 2,5-dihydroxybenzoic acid was prepared as a 20 mg/mL solution is 2:1 CHCl_3_:MeOH (v/v) and applied to the sample using a SunCollect automatic pneumatic sprayer (Sunchrom GmbH, Friedrichsdorf, Germany). In total 15 layers of matrix were deposited onto the tissue. The first three layers were deposited using a flow rate of 10, 20 and 30 μL/min, respectively, and all subsequent layers at 40 μL/min.

### Haematoxylin and eosin (H&E) staining

Haematoxylin and eosin (H&E) staining was performed on tissue sections post-imaging. The matrix was removed by immersion in 100% methanol for 30 s, after which the following protocol was used. The tissues were first washed with series of solutions (2 × 95% EtOH, 2 × 70% EtOH and deionised water for 2 min each), stained with haematoxylin for 3 min and subsequently washed with running tap water for 3 min. Tissues were then stained with eosin for 30 s and washed with running tap water for 3 min. After staining, the samples were placed into 100% ethanol for 1 min, then in xylene for 30 s. Finally, glass coverslips were placed onto the samples using Entellen mounting medium. High-resolution optical scans of the stained tissue sections were acquired using a Leica Aperio CS2 (Leica Biosystems Imaging, Vista, CA, USA).

### MALDI-MSI

MALDI-MSI was performed using a hybrid Orbitrap Elite mass spectrometer (Thermo Fisher Scientific GmbH, Bremen, Germany) coupled to a reduced pressure ESI/MALDI ion source (Spectroglyph LLC, Kennewick, WA, USA). Further details on the experimental setup can be found in reference^38^. All MSI data were collected using a nominal mass resolution setting of 240,000 @ *m/z* 400, an *m/z* range of 350–1000 and a pixel size of 40 × 40 μm^2^. Tandem mass spectrometry (MS/MS) was performed on MALDI ions generated directly from lung tissue sections to confirm the identification of the ions of interest. MS/MS spectra were acquired using resonant collision-induced dissociation, a ± 0.5 Da isolation window and Orbitrap detection of fragments.

### Data analysis

MSI data analysis and visualisation were performed using LipostarMSI (Molecular Horizon Srl, Perugia, Italy)^39^. During data import, mass spectra were recalibrated using the [PC32:0+Na]^+^ ion as a lock mass (theoretical *m/z* 756.551374). Peak picking was performed using a relative intensity threshold of 0.05%, a minimum peak frequency of 0.5% and an *m/z* alignment tolerance of 3 ppm. All MSI data visualisations were generated using total ion current normalisation with hotspot removal (quantile thresholding, high quantile = 99%). An exception was the ratio images that were produced by normalisation to the denominator ion signal ([PC32:0+Na]^+^ or ([PC32:1+Na]^+^.

Xcalibur QualBrowser (version 4.1.50, Thermo Fisher Scientific GmbH, Bremen, Germany) and Python 3.7.6 together with the pymsfilereader 1.0.1 module (https://pypi.org/project/pymsfilereader/) were used for analysis and visualisation of .raw MS and MS/MS data spectral data.

### Lipid nomenclature

Sum-composition PC molecular species are generally denoted by the number of total fatty acyl carbon atoms followed by the number of unsaturated double bonds. For example, PC36:4 corresponds to a PC lipids with 36 carbons and 4 double bonds distributed across the two acyl chains. In cases where the length and degree of unsaturation of each acyl chain are known but the *sn*-positioning of individual fatty acyls is undefined, the underscore nomenclature is used, for example PC16:0_20:4. When the *sn*-positioning is defined, the “/” nomenclature is used. For example PC16:0/20:4 corresponds to a PC lipid with a 16:0 acyl chain at the *sn*-1 position and a 20:4 acyl chain at the *sn*-2 position. Stereochemistry and double bond positions are undefined. Sum-composition lipid species can consist of multiple molecular species. For instance, PC36:4 could be either PC18:2_18:2 or PC PC16:0_20:4. The exception is for newly synthesised PC species containing ^13^C_24_-palmitoyl-lysoPC derived from U^13^C-DPPC metabolism where, for instance, ^13^C_24_-PC36:4 is unambiguously the palmitoylarachidonoyl PC16:0_20:4 species.

## Acknowledgements

This work was financially supported by the LINK program of the Dutch province of Limburg. SRE acknowledges funding from the Australian Research Council Future Fellowship Scheme (grant number FT190100082) and the Netherlands Organisation for Scientific Research VIDI scheme (grant number 198.011). We are grateful to the NIHR Southampton Biomedical Research Centre for mass spectrometry support and to Chiesi Farmaceutica for the supply of the therapeutic surfactants.

## Data availability statement

Mass spectrometry imaging datasets in the open source imzML format are available for download from the MetaboLights repository^40^ with study identifier MTBLS2075 (www.ebi.ac.uk/metabolights/MTBLS2075).

## Author Contributions

SRE, RMAH and ADP conceived the project. SRE and BF performed MSI sample preparation and data acquisition. HWC and JM deigned the mouse experiments and performed the mouse procedures, EH and MP performed the alveolar macrophage procedures and GK and MP provided mass spectrometry. SRE and ADP analysed the data. SRE, RMAH, ADP, JM and HWC financially supported the project. SRE and ADP wrote the paper with input from co-authors.

