## Supporting Information for "Mass spectrometry imaging of phosphatidylcholine metabolism in lungs administered with therapeutic surfactants and isotopic tracers"

*To whom correspondence should be addressed:

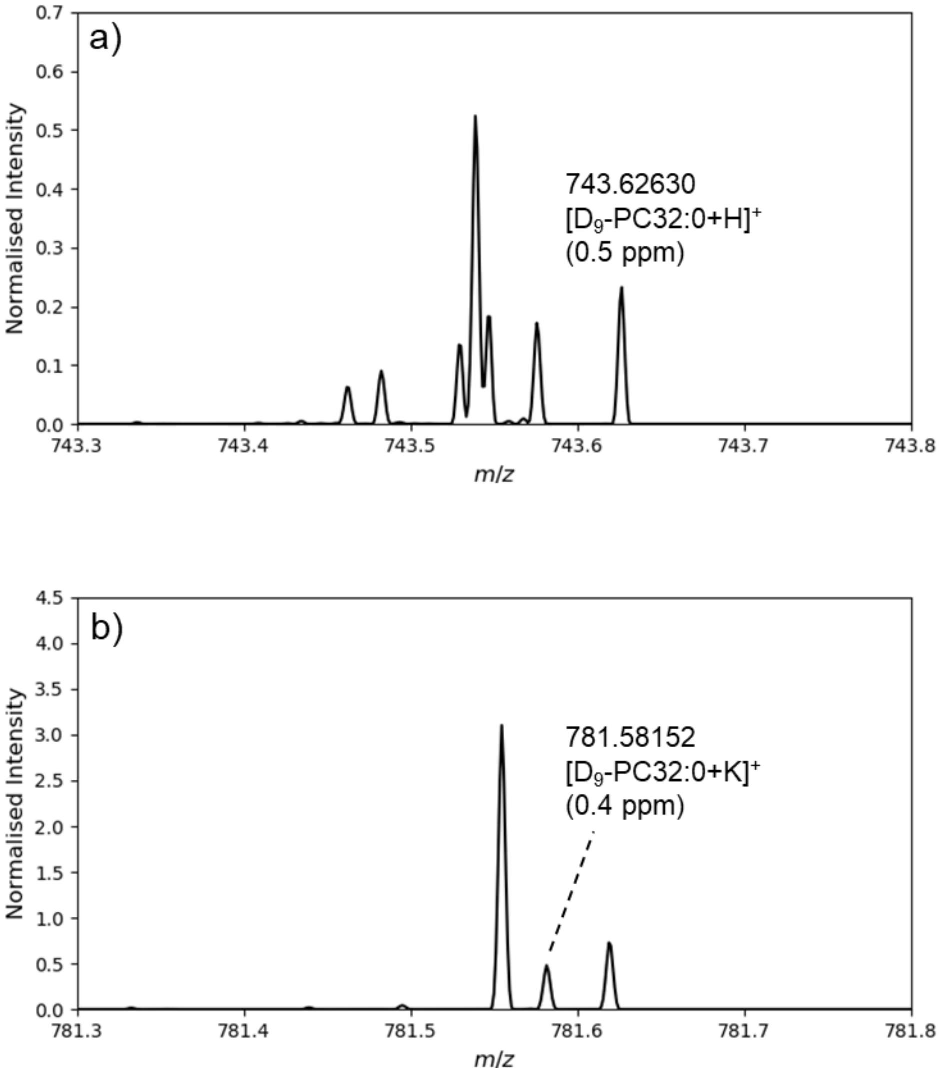


**Figure S1.** Zoomed-in regions of the averaged positive-ion MALDI spectrum acquired from mouse lung tissue dosed with $D_9$-choline and U-$C_{13}$-DPPC–containing CHF5633 surfactant (labels administered 12 h prior to sacrifice). Spectra demonstrate the detection of (a) ([$D_9$-PC(32:0)+H]$^+$ and (b) ([$D_9$-PC(32:0)+K]$^+$). Parts-per-million (ppm) mass errors are indicated in parentheses and spectra are normalised to the base peak intensity (0–100%).

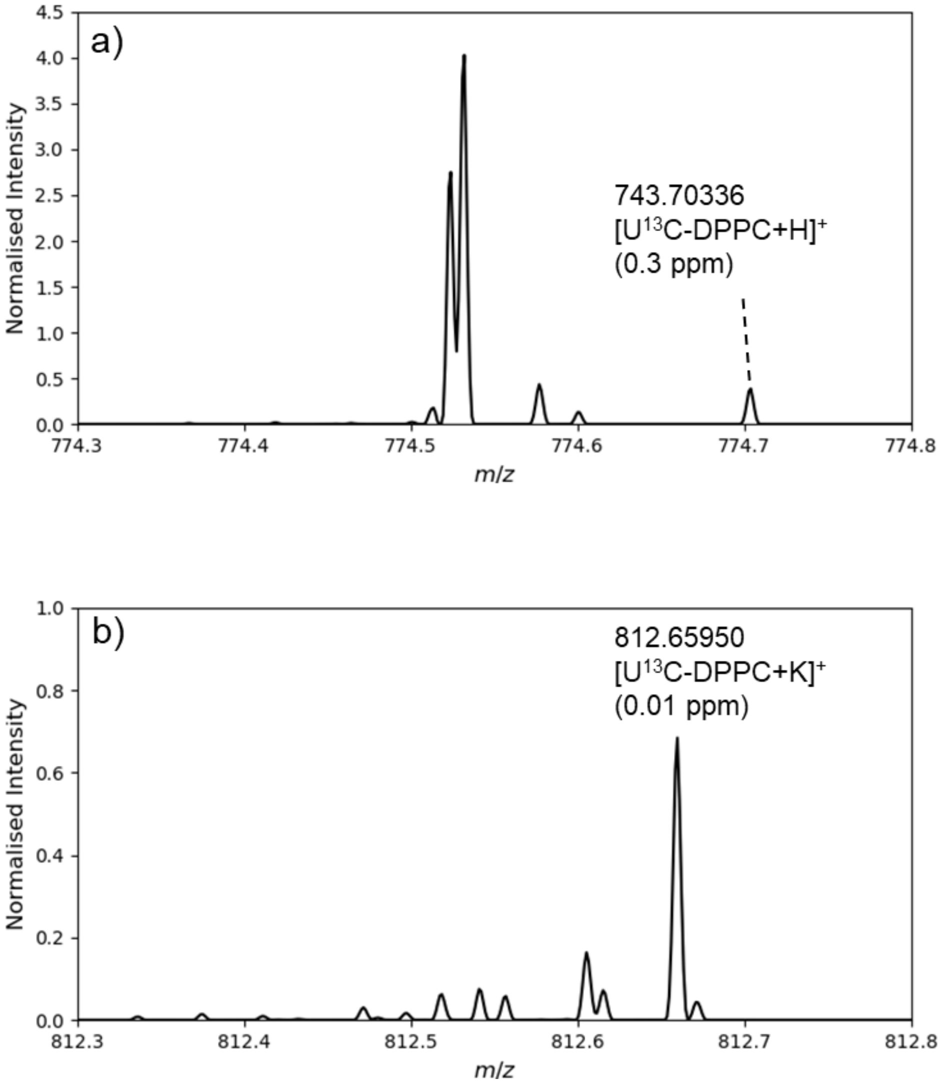


**Figure S2.** Zoomed-in regions of the averaged positive-ion MALDI spectrum acquired from mouse lung tissue dosed with $D_9$-choline and U$^{13}$C-DPPC–containing CHF5633 surfactant (labels administered 12 h prior to sacrifice). Spectra demonstrate the detection of (a) ([U$^{13}$C-DPPC+H]$^{+}$ and (b) ([U$^{13}$C-DPPC+K]$^{+}$). Parts-per-million (ppm) mass errors are indicated in parentheses and spectra are normalised to the base peak intensity (0–100%). DPPC = PC16:0/16:0

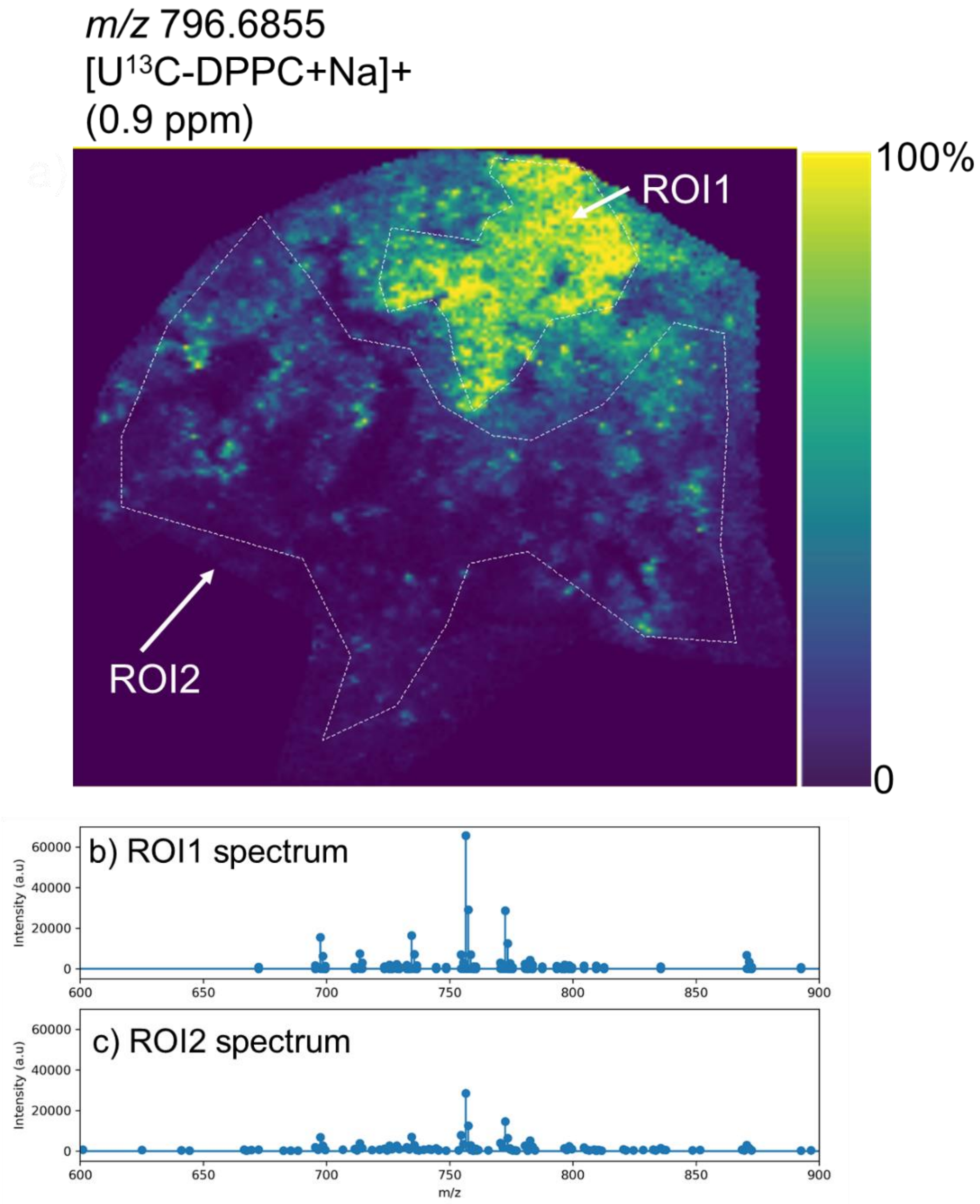


**Figure S3.** (a) Ion distribution image of $[U^{13}C\text{-}DPPC+Na]^+$ observed at *m/z* 796.6855 in mouse lung tissue 12 h after nasal administration of CHF5633 surfactant. Signal for $[U^{13}C\text{-}DPPC+Na]^+$ is specific for CHF5633. (b, c) Extracted region-of-interest (ROI) spectra from the corresponding regions marked in (a) demonstrating the overall increased signal of unlabelled $[PC32{:}0+Na]^+$ in regions of CHF5633 accumulation.

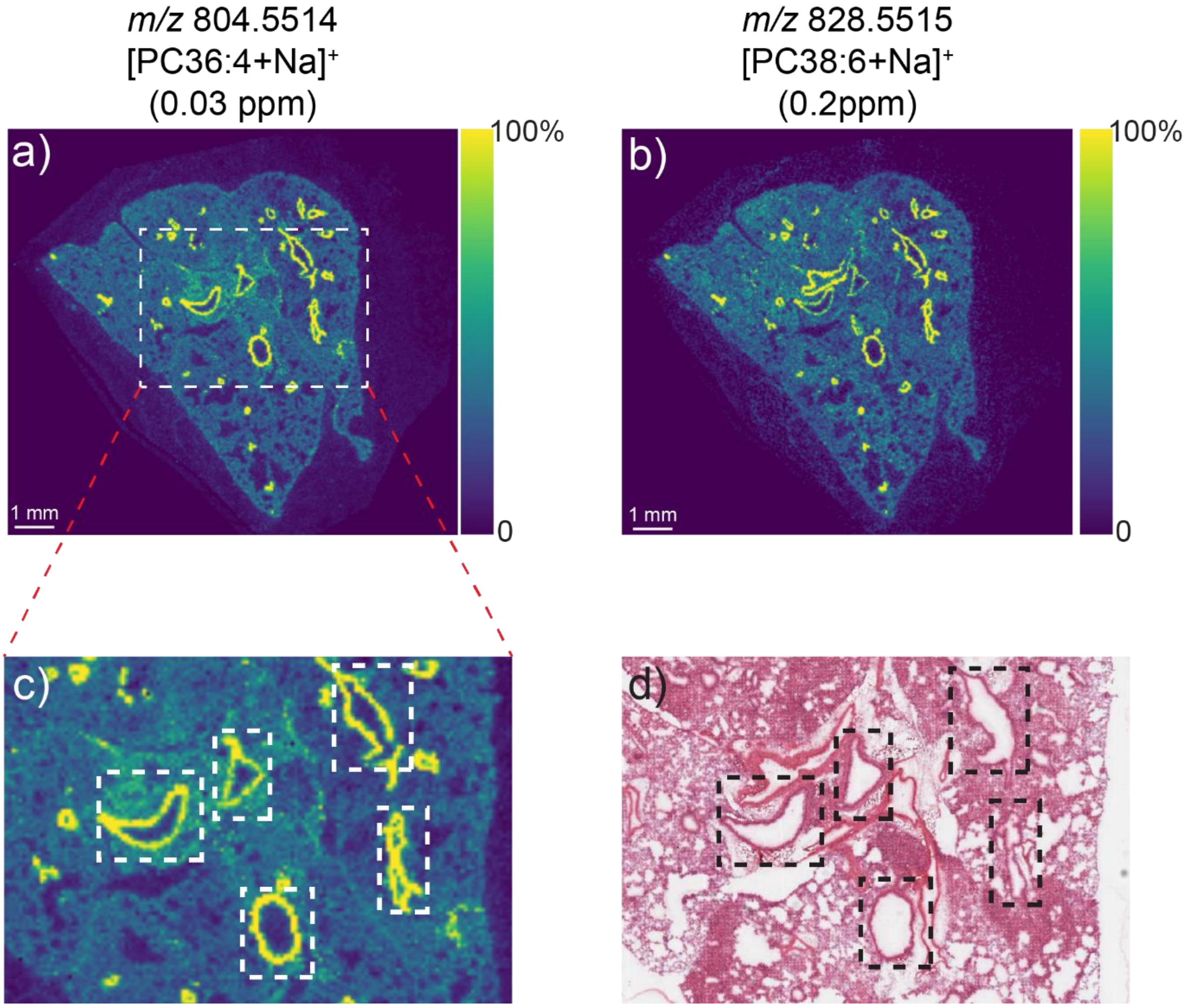


**Figure S4.** Ion distribution images for (a) $[PC36:4+Na]^+$ (*m/z* 804.5514) and (b) $[PC38:6+Na]^+$ (*m/z* 828.5515) obtained from mouse lung tissue collected 6 h after administration of $D_9$-choline and $U^{13}C$-DPPC–containing CHF5633. Parts-per-million (ppm) mass errors are indicated in parentheses. (c) Magnification of the boxed region in (a) with selected bronchiolar regions outlined in white boxes. (d) The corresponding H&E-stained tissue section with the same selected bronchiolar regions outlined in black boxes. These data demonstrate the co-localisation of the polyunsaturated lipids PC36:4 and PC38:6 with the bronchiolar regions of the lung. All MSI images were visualised using total ion current normalisation and hotspot removal (high quantile = 99%).

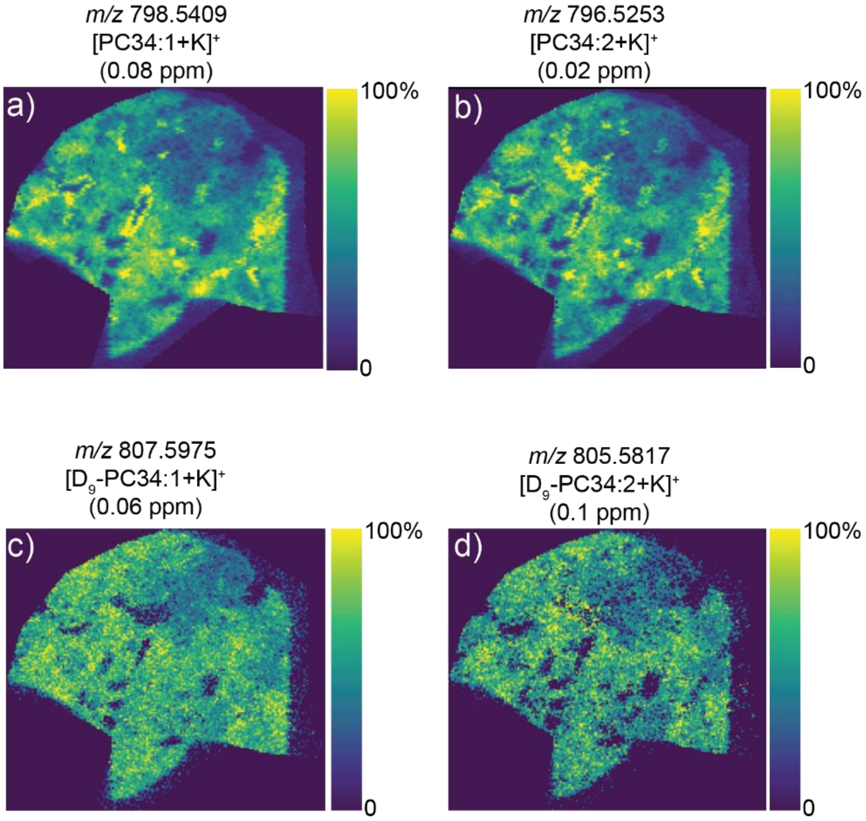


**Figure S5.** Ion distribution images for (a) $[PC34:1+K]^+$ (*m/z* 798.5409), (b) $[PC34:2+K]^+$ (*m/z* 796.5253), (c) $[D_9\text{-}PC34:1+K]^+$ (*m/z* 807.5975) and (d) $[D_9\text{-}PC34:2+K]^+$ (*m/z* 805.5817) obtained from the same tissue as that shown in Figures 2 and 3 in the main text. Parts-per-million (ppm) mass errors are indicated in parentheses. All MSI images were visualised using total ion current normalisation and hotspot removal (high quantile = 99%).

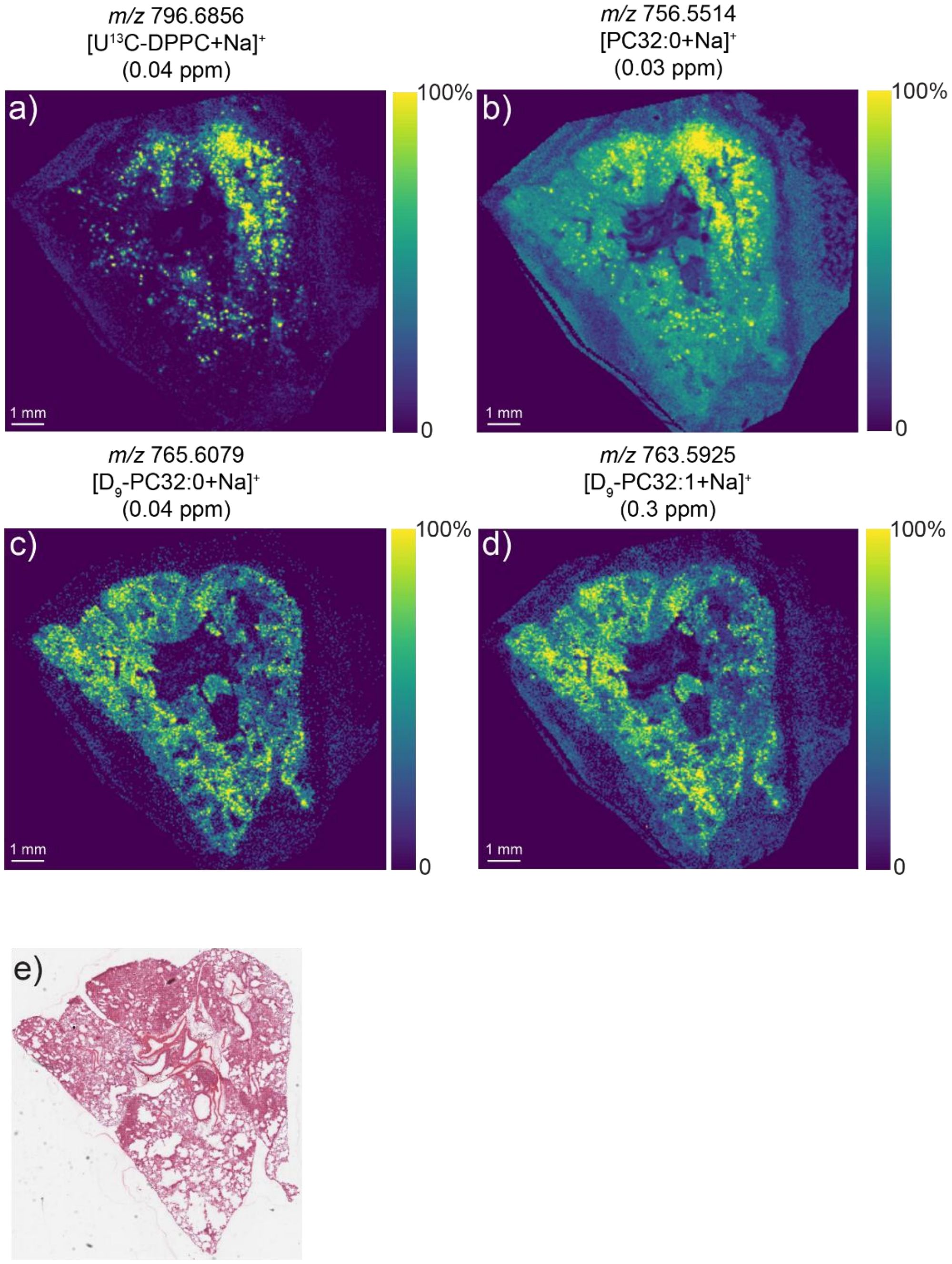


**Figure S6.** MALDI-MSI data from mouse lung tissue administered with $D_9$choline and $U^{13}C$-DPPC–containing CHF5633 surfactant (labels administered 6 h prior to sacrifice, same tissue as shown in Figure S4). (a–d) Ion images of (a) *m/z* 796.6855 ([$U^{13}C$-DPPC+Na]$^+$), (b) *m/z* 756.5514 ([PC32:0+Na]$^+$, (c) *m/z* 765.6079 ([$D_9$-PC32:0+Na]$^+$) and (d) *m/z* 754.5359 ([PC32:1+Na]$^+$). (e) Optical image of post-MSI H&E-stained tissue section. Part-per-million (ppm) mass errors are indicated in parentheses. All images were visualised using total-ion-current normalisation and hotspot removal (high quantile = 99%). DPPC = PC16:0/16:0.

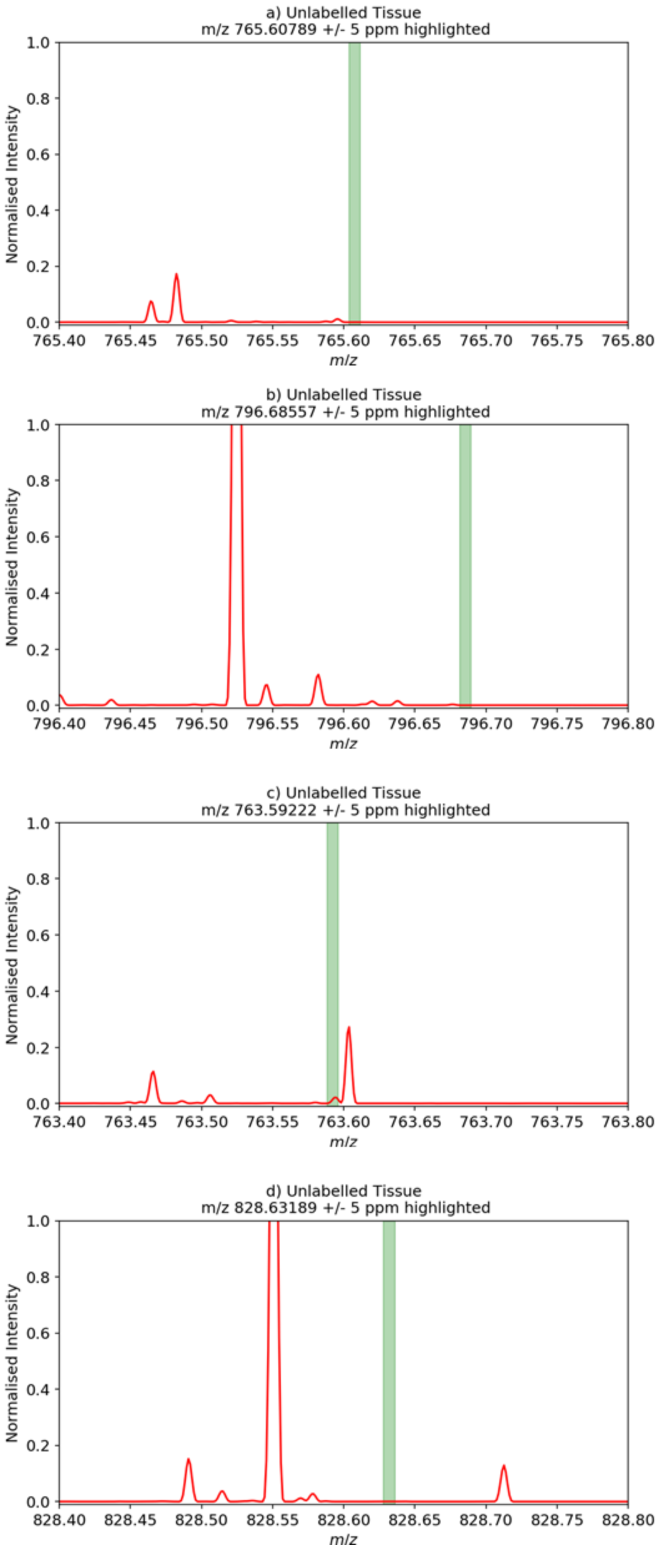


**Figure S7.** Averaged MALDI mass spectra obtained from control mouse lung tissue not administered with either $D_9$-choline or $U^{13}C$-DPPC. The expected *m/z* ranges (± 5 ppm) where signal for (a) $[D_9\text{-PC32:0+Na}]^+$, (b) $[U^{13}\text{-DPPC+Na}]^+$, (c) $[D_9\text{-PC32:1+Na}]^+$, and (d) $[^{13}C_{24}\text{-PC16:0\_20:4+Na}]^+$ is expected are indicated in green. Spectra are normalised to the base peak intensity (0–100%). DPPC = PC16:0/16:0.

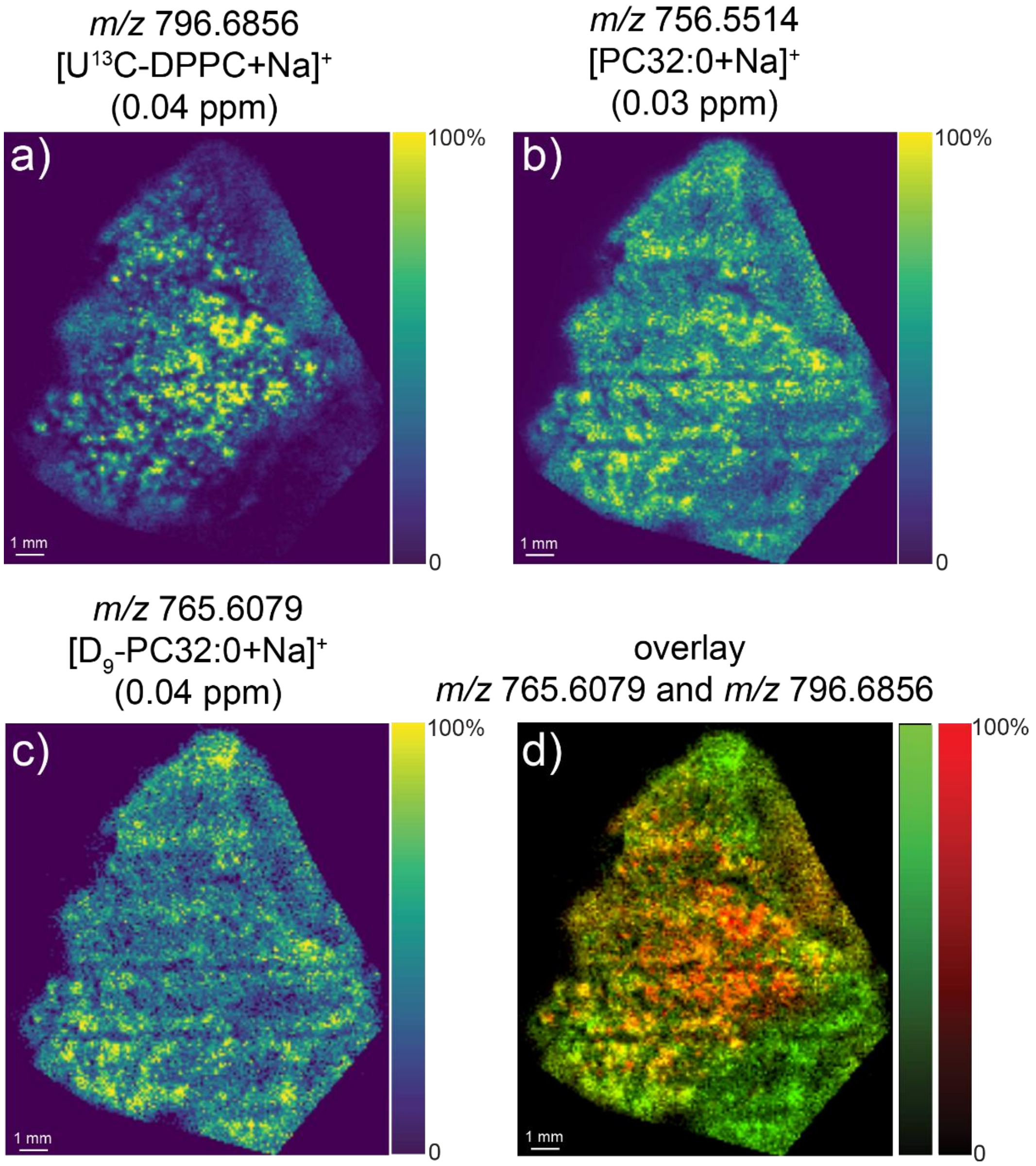


**Figure S8.** MALDI-MSI data of mouse lung tissue administered with $D_9$-choline and U $^{13}C$-DPPC–containing Poractant alfa surfactant (labels administered 18 h prior to sacrifice). Ion images of (a) *m/z* 796.6856 ([U$^{13}$C-DPPC+Na]$^+$), (b) *m/z* 756.5154 [PC32:0+Na]$^+$ and (c) *m/z* 765.6079 ([$D_9$-PC32:0+Na]$^+$). (d) Overlay image of $D_9$-PC32:0+Na]$^+$ (red) and [U-$^{13}$C-DPPC+Na]$^+$ (green). Parts per million (ppm) mass errors are indicated in parentheses. All images were visualised using total-ion-current normalisation and using hotspot removal (high quantile = 99%). DPPC = PC16:0/16:0.

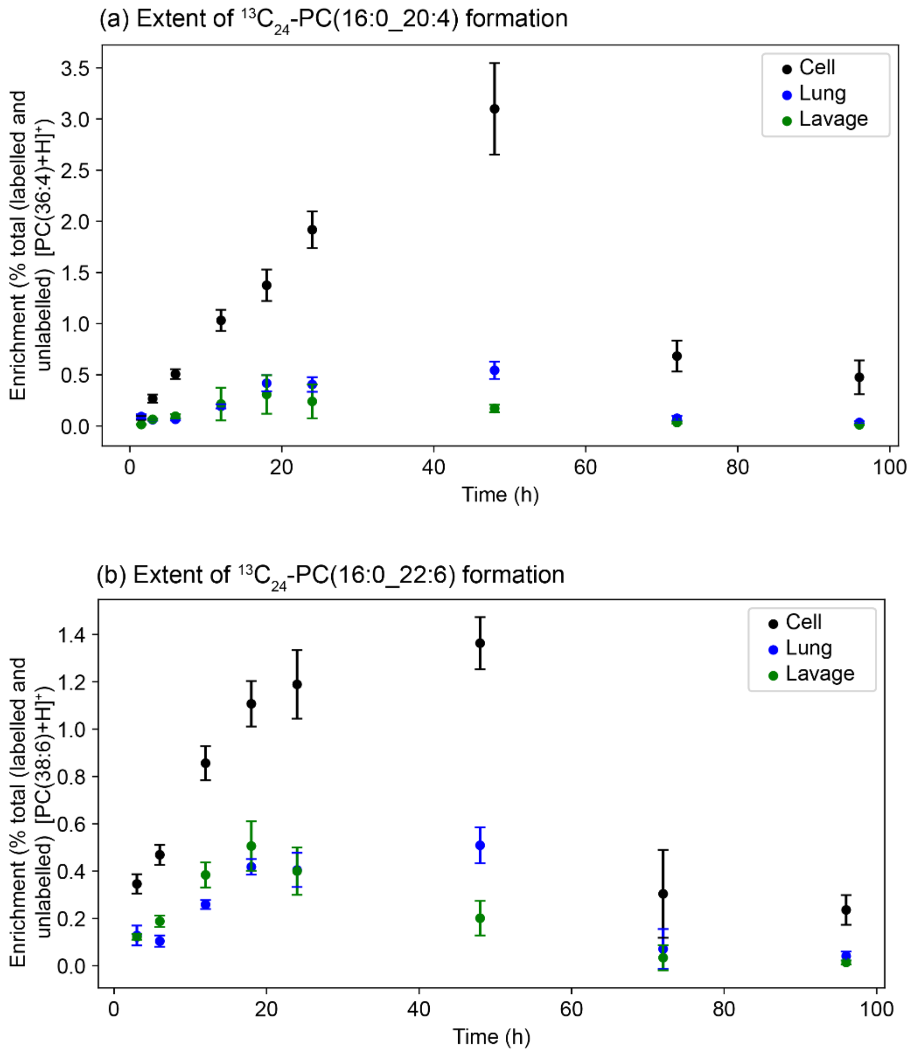


**Figure S9.** ESI-MS analysis of lavage cell pellet, lung tissue and lung lavage lipid extracts collected from animals ($n$ = 10–21 mice per time point/group) at the indicated time after nasal administration of U-$^{13}$C-DPPC–containing Poractant alfa surfactant. The extent of formation of acyl remodelling products (a) $^{13}C_{24}$-PC16:0/20:4 and (b) $^{13}C_{24}$-PC16:0/22:6 originating from acyl remodelling of U$^{13}$C-DPPC was followed over 96 hours. Error bars represent ± 1 standard deviation. $^{13}C_{24}$-labelled lipid species were detected using a precursor ion scan of $m/z$ 189.0 for the $^{13}C_5$-labelled phosphocholine headgroup. Presented data are a subsequent analysis of data collected in reference [12]. There was a high variability in nasal delivery of surfactant to the mice in [12], with some mice receiving little surfactant. As the purpose of this figure was to present a comparison between matched sub-fractions of lung rather than comparing absolute concentrations, mice with a low abundance of delivered surfactant were excluded from this comparison, although reported in reference 12. The criterion used was an enrichment of exogenous surfactant < 0.1 % of total BALF phosphatidylcholine, based on the analysis of U$^{13}$C-DPPC.

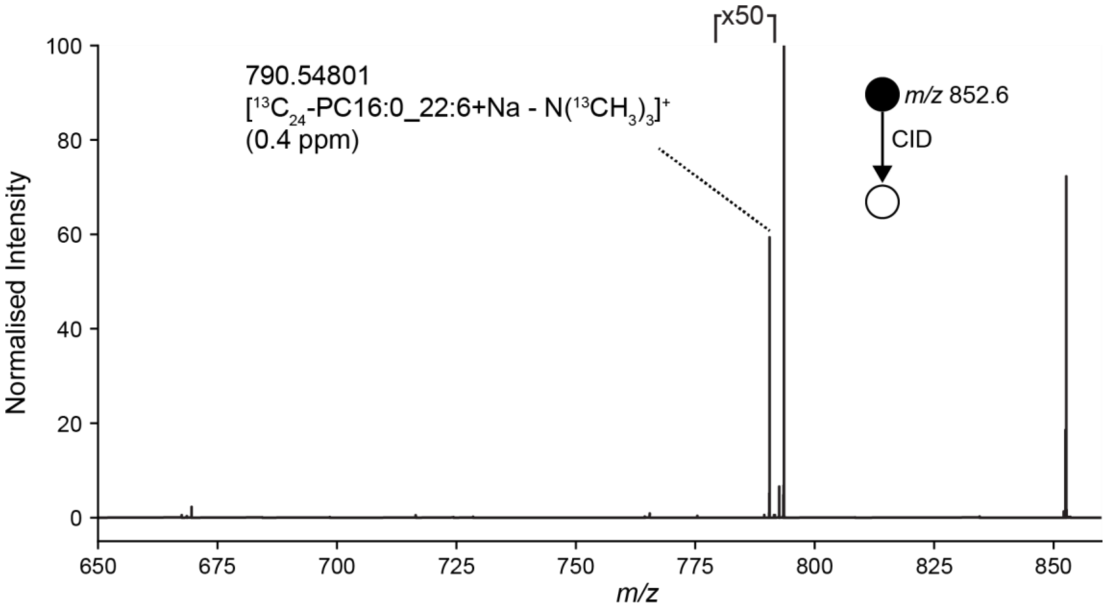


**Figure S10.** MS/MS spectrum of precursor ions at *m/z* 852.6 ± 0.5 with fragment ions consistent with the presence $[^{13}C_{24}$-PC16:0_22:6+Na$]^+$ annotated. Parts per million (ppm) mass errors are provided in parentheses.
